# Mucociliary Transport Deficiency and Disease Progression in Syrian Hamsters with SARS-CoV-2 Infection

**DOI:** 10.1101/2022.01.16.476016

**Authors:** Qian Li, Kadambari Vijaykumar, Scott E Philips, Shah S Hussain, Van N Huynh, Courtney M Fernandez-Petty, Jacelyn E Peabody Lever, Jeremy B Foote, Janna Ren, Javier Campos-Gómez, Farah Abou Daya, Nathaniel W Hubbs, Harrison Kim, Ezinwanne Onuoha, Evan R Boitet, Lianwu Fu, Hui Min Leung, Linhui Yu, Thomas W Detchemendy, Levi T Schaefers, Jennifer L Tipper, Lloyd J Edwards, Sixto M Leal, Kevin S Harrod, Guillermo J Tearney, Steven M Rowe

**Affiliations:** Departments of Medicine, University of Alabama at Birmingham, Birmingham, AL, United States; Departments of Gregory Fleming James Cystic Fibrosis Research Center, University of Alabama at Birmingham, Birmingham, AL, United States; Departments of Graduate Biomedical Sciences Program, University of Alabama at Birmingham, Birmingham, AL, United States; Departments of Microbiology, University of Alabama at Birmingham, Birmingham, AL, United States; Departments of Chemistry, University of Alabama at Birmingham, Birmingham, AL, United States; Departments of Radiology, University of Alabama at Birmingham, Birmingham, AL, United States; Departments of Biomedical Engineering, University of Alabama at Birmingham, Birmingham, AL, United States; Wellman Center for Photomedicine, Massachusetts General Hospital, Harvard Medical School, Boston, MA, United States; Departments of Anesthesiology and Perioperative Medicine, University of Alabama at Birmingham, Birmingham, AL, United States; Departments of Biostatistics, University of Alabama at Birmingham, Birmingham, AL, United States; Departments of Pediatrics, University of Alabama at Birmingham, Birmingham, AL, United States; Departments of Cell Developmental and Integrative Biology, University of Alabama at Birmingham, Birmingham, AL, United States

## Abstract

Substantial clinical evidence supports the notion that ciliary function in the airways plays an important role in COVID-19 pathogenesis. Although ciliary damage has been observed in both *in vitro* and *in vivo* models, consequent impaired mucociliary transport (MCT) remains unknown for the intact MCT apparatus from an *in vivo* model of disease. Using golden Syrian hamsters, a common animal model that recapitulates human COVID-19, we quantitatively followed the time course of physiological, virological, and pathological changes upon SARS-CoV-2 infection, as well as the deficiency of the MCT apparatus using micro-optical coherence tomography, a novel method to visualize and simultaneously quantitate multiple aspects of the functional microanatomy of intact airways. Corresponding to progressive weight loss up to 7 days post-infection (dpi), viral detection and histopathological analysis in both the trachea and lung revealed steadily descending infection from the upper airways, as the main target of viral invasion, to lower airways and parenchymal lung, which are likely injured through indirect mechanisms. SARS-CoV-2 infection caused a 67% decrease in MCT rate as early as 2 dpi, largely due to diminished motile ciliation coverage, but not airway surface liquid depth, periciliary liquid depth, or cilia beat frequency of residual motile cilia. Further analysis indicated that the fewer motile cilia combined with abnormal ciliary motion of residual cilia contributed to the delayed MCT. The time course of physiological, virological, and pathological progression suggest that functional deficits of the MCT apparatus predispose to COVID-19 pathogenesis by extending viral retention and may be a risk factor for secondary infection. As a consequence, therapies directed towards the MCT apparatus deserve further investigation as a treatment modality.

## Introduction

Coronavirus disease 2019 (COVID-19), caused by severe acute respiratory syndrome coronavirus 2 (SARS-CoV-2), is dominated by respiratory disease including pneumonia^1–4^. SARS-CoV-2 binds to its cellular receptor, angiotensin-converting enzyme 2 (ACE2), initiating cell entry and subsequent pathogenesis^5–9^. The degree of ACE2 expression is thought to impart susceptibility to SARS-CoV-2 and thus the severity of the disease upon infection^10–13^. Single-cell RNA sequencing (scRNA-seq) from healthy donors has demonstrated high expression of ACE2 in respiratory epithelial cells^14^, indicating tropism of SARS-CoV-2 infection for the respiratory tract^15–17^. Epithelial cells represent the first contact of respiratory viruses to the host, followed by pathological progression to the lung and other tissues as descending infection occurs. Thus, the functional consequences of epithelial cell infection in COVID-19 are crucial to understanding its pathogenesis.

Among respiratory epithelial cells, ACE2 expression dominates in ciliated cells^14^, a cell type found throughout the conducting airways and crucial for maintaining mucociliary transport (MCT), a critical host defense mechanism. Excessive mucin production and hyperviscous mucus have been frequently found in bronchoalveolar lavage fluid (BALF)^18–20^ and airways^21,22^ of COVID-19 patients. Perhaps consequently, bacterial or fungal superinfection occurs in up to one-third of COVID-19 patients^23–26^ and contributes to increased severity and mortality^24–27^. *In vitro* studies using differentiated human tracheobronchial epithelial (HTBE) cells showed SARS-CoV-2 infection shrinks cilia and abolished ciliary beating in the center of cytopathic plaques, with additional morphologic changes to cilia in neighboring regions^28^. Impaired MCT was reported in HTBE cells measured by tracking polystyrene beads^29^. These reports suggest that diminished MCT could contribute to descending viral infection and secondary infections, but presently remains unknown for the intact mucociliary clearance (MCC) apparatus (i.e., tissue level analysis) from an *in vivo* model of disease.

Syrian hamsters have been used to model respiratory infections by SARS-CoV, influenza virus, and adenovirus^30–33^. ACE2 sequence homology at the SARS-CoV-2 receptor binding domain has shown hamsters to have the most similarity to humans, second only to non-human primates^34^. Furthermore, hamsters with SARS-CoV-2 exhibit virological, physiological, and pathological endpoints relevant to human disease^35–38^, being applied to various studies including mechanistic investigation^34,39–43^, vaccine^44–48^, antibody^49,50^, and therapeutic^51^ development. Here, we followed the time courses of virological, physiological, and pathological changes in hamsters up to 7 days post-infection (dpi) of SARS-CoV-2, and monitored changes to the MCT apparatus using micro-optical coherence tomography (μOCT), a novel method to visualize and simultaneously quantitate multiple aspects of the functional microanatomy of airways in intact tissues^52^.

## Results

### SARS-CoV-2 induces weight loss and lethargy in hamsters

To assess the effects of SARS-CoV-2 infection on the mucociliary transport apparatus, we first established the model in Golden Syrian Hamsters, an animal model that recapitulates human COVID-19^35–38^. Under BSL-3 conditions and as shown in Fig. 1, following nasal inoculation of 3 × 10^5^ plaque-forming units (PFU) of SARS-CoV-2 in 100 μL of viral propagation media, hamsters exhibited progressive weight loss up to 10.2 ± 1.0% of body weight, peaking at 6 dpi (Fig. 1, P=0.0052) as compared to stable body weight observed in mock controls evaluated by the same experimental procedures. This is consistent with previous hamster studies using various strains and titers of SARS-CoV-2 that demonstrated peak bodyweight loss that ranged from 10–20% and peaked between 5 and 7 dpi^34,37–39,41,42,44,46,53^. We also observed moderate lethargy through 7 dpi in SARS-CoV-2 infected hamsters but not in mock controls, although other clinical signs such as changes of fur or posture, sneezing, coughing, diarrhea, and nasal or ocular discharge were not found. This clinical syndrome was also consistent with previous reports using other strains of SARS-CoV-2 in hamsters^34,36^.

**Figure 1.**
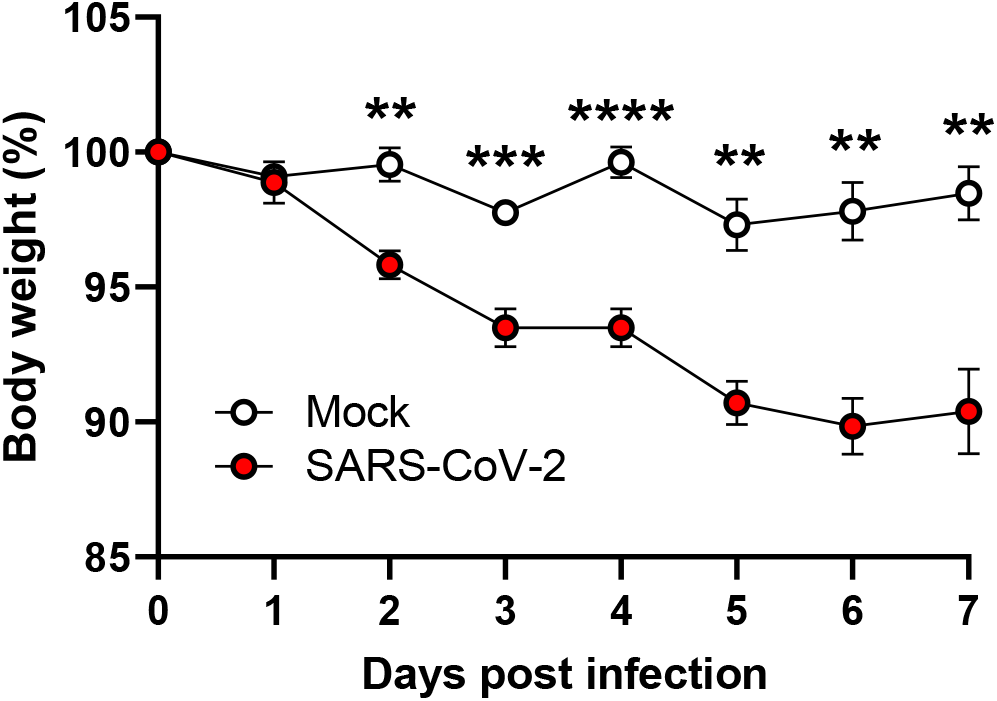
Reduced body weight in hamsters through 7 days post-infection with SARS-CoV-2. Golden Syrian hamsters were inoculated intranasally with 3×10^5^ plaque-forming units of SARS-CoV-2 or vehicle (mock), and body weight was monitored up to 7 days post-infection (dpi; N=8-25 for infected and N=4-12 for mock). P<0.0001 for time, infection, and interaction by two-way ANOVA; **P<0.01, ***P<0.001 and ****P<0.0001 by Šídák’s posthoc test.

### SARS-CoV-2 is prominently located in the respiratory tract

To evaluate viral infection, we detected SARS-CoV-2 in hamsters using complementary techniques. Figure 2A shows viral load in various samples at 2, 4, and 7 dpi quantified by quantitative real-time reverse transcriptase-polymerase chain reaction (qRT^2^-PCR). We detected high viral titers in the respiratory tract that exceeded 10^7^ genome copies/ml through 4 dpi, including nasal brush, nasal wash, and BALF specimen. Viral titers in these respiratory samples were notably greater than titers in either the gastrointestinal (GI) tract (oral and rectal swabs were approximately 10^5^ and 10^3^ copies/ml, respectively) or in circulation (*i.e*., serum, approximately 10^2^ copies/ml). Although differences in sample collection methods or sample volume may have contributed to the differences observed, viral titers in the respiratory tract exceeded other tissues by at least 3-orders of magnitude. This indicated important differences among these tissues that make the respiratory tract the main target of viral invasion, which is consistent with the tissue specificity reported by Chan *et al*.^34^. Higher viral load in BALF than in oral and anal swabs has also been found in COVID-19 patients^9^.

**Figure 2.**
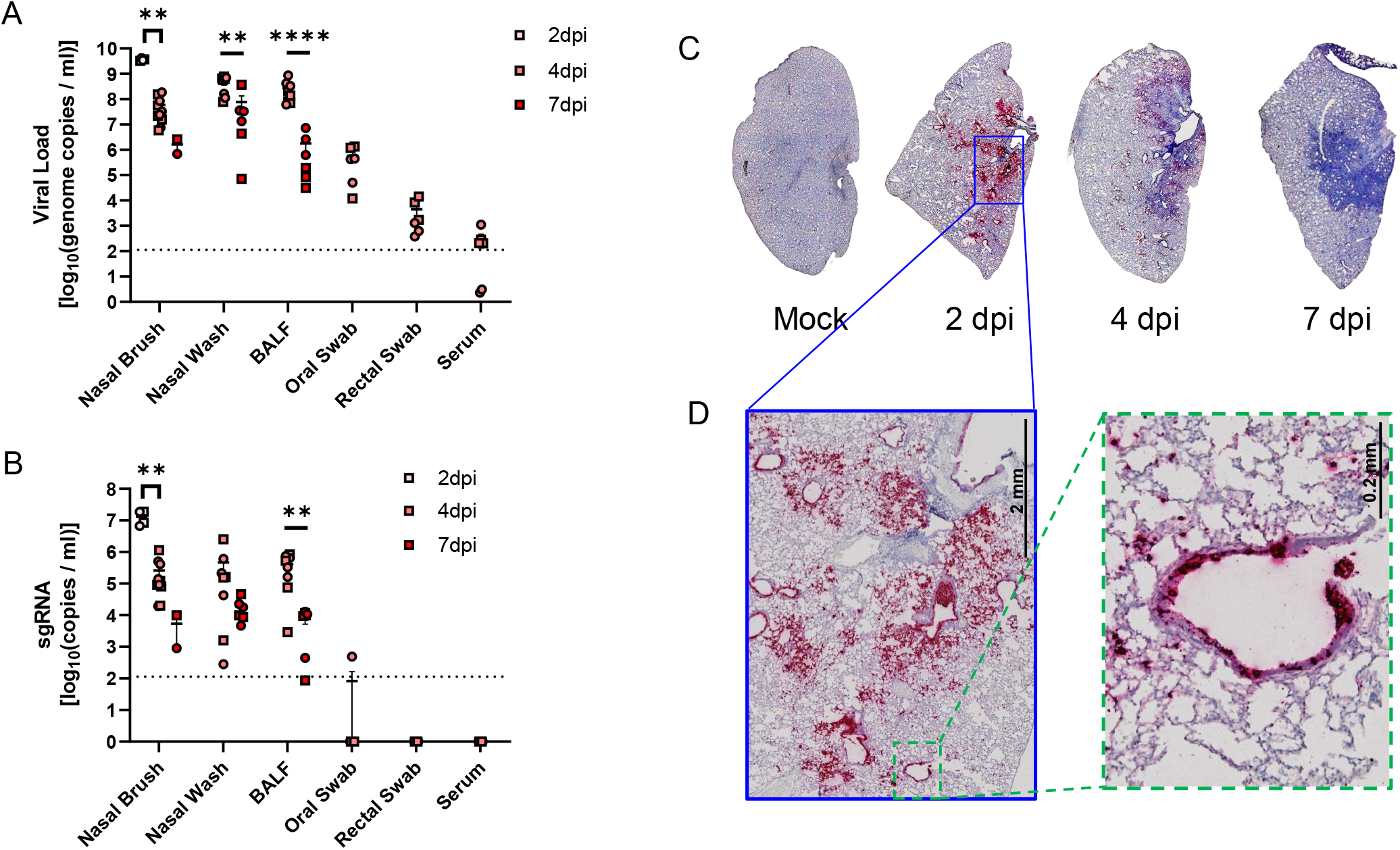
SARS-CoV-2 detection in hamsters through 7 days post-infection. Golden Syrian hamsters were inoculated in Fig.1. Then samples were collected at 2, 4, and 7 dpi. Q-RT^2^-PCR vs. standard curve quantitated genomic (A) and subgenomic (B) viral titers for nasal brush, nasal wash, bronchial alveolar lavage fluid (BALF), oral swab, rectal swab, and serum. Viral titers in all types of samples from mock were under the detection limit, therefore, not shown. Subgenomic viral titers in serum, rectal swab, and most oral swabs were below quantitation limits by PCR, and are shown as 0. The dotted line indicates the limits of the PCR method. Square represents male and circle female. Values in log_10_(copies/ml) were used for statistical analysis. For nasal brush, P<0.0001 and P=0.0161 for genomic and subgenomic titers, respectively, by one-way ANOVA with **P<0.01 (shown by a bracket) by following Tukey’s posthoc test. For nasal wash and BALF, **P<0.01 and ****P<0.0001 (shown by straight lines) by unpaired t-test. Representative images of the whole left lobe slices with SARS-CoV-2 detection by RNAscope^®^ (C), with high power view of the airway (left) and parenchymal (right) at 2 dpi (D).

We also performed analysis using primers to detect subgenomic (sg) SARS-CoV-2 RNA by qRT^2^-PCR to confirm SARS-CoV-2 replication. Along with the analysis of genomic copies, the respiratory tract was the dominant site of viral replication (Fig. 2B). At 4 dpi, sgRNA levels exceeded 10^5^ copies/ml in the nasal brush, nasal wash, and BALF specimens, whereas in oral, rectal swabs, and sera, those were below quantitation limits.

The viral load decreased with time by examining differences in viral titers of the respiratory tract within the same sample type (Fig. 2A). In nasal brush specimens, the viral RNA copy number at 2, 4, and 7 dpi decreased significantly from 9.6±0.0 to 7.5±0.2 and then 6.1±0.3 log_10_(copies/ml), respectively (P<0.0001). For nasal wash and BALF, similar findings were reported at 4 and 7 dpi, with a reduction of 8.5±0.1 to 7.0±0.5 log_10_(copies/ml) for nasal wash (P=0.0077), and 8.3±0.1 to 5.6±0.4 log_10_(copies/ml) for BALF(P<0.0001). Levels of sgRNA detected within a sample type in the respiratory tract also diminished with time (7.1±0.1 *vs*. 5.1±0.2 *vs*. 3.5±0.5 log_10_(copies/ml) at 2, 4, and 7 dpi for nasal brush, P=0.0161; 5.3±0.3 *vs*. 3.5±0.4 log_10_(copies/ml) for BALF at 4 and 7 dpi, P=0.0021). High viral loads at the onset of infection that diminished over time are consistent with the previous reports^34,38–40,46,53^.

Next, we examined SARS-CoV-2 RNA distribution in hamster lungs by *in situ* hybridization with RNAscope^®^. Each viral RNA transcript was hybridized and then the detection signal was visualized as a point of amplification. Figures 2C and 2D show that SARS-CoV-2 in the lung exhibited an airway-centric pattern. Large and medium airways exhibited the most prominent viral RNA detection, whereas parenchymal distribution was patchy and heterogeneous (Fig. 2D). This pattern evolved with peak detection at 2 dpi that remained prominent and airway centric at 4 dpi, but was absent by 7 dpi (Fig. 2C). Together with genomic and subgenomic viral RNA found in respiratory samples by qRT^2^-PCR, our results demonstrate prominent viral RNA presence in the respiratory airways that is most evident in the large and medium airways and progresses to the distal lung thereafter. The data further show clearance of SARS-CoV-2 at 7 dpi, which is consistent with the recovery and weight gain after 6 or 7 dpi. These findings indicate respiratory epithelial cells as the main target of initial viral infection.

### SARS-CoV-2 infection causes lung injury in hamsters

The major focus of severe COVID-19 has been the injury to the distal lung parenchyma; accordingly, we evaluated lesions in the lung to characterize the pathology induced by SARS-CoV-2 infection in hamsters. As SARS-CoV-2 was gradually depleted, the lung damage caused by initial viral invasion progressed with time and increased in severity through 7 dpi, in concert with changes in viral replication (Fig. 3). Lung to body weight (L/BW) ratio (Fig. 3A) steadily increased by 62% from 4.7±0.1 in mock to 7.6±0.7 mg/g 7 dpi (P<0.0001), suggesting progressive development of edema and cell infiltration. Multifocal consolidation and hyperemia were apparent as early as 4 dpi and were greatest at 7 dpi on gross inspection (Fig. 3B). The hematoxylin and eosin (H&E) stained lungs of SARS-CoV-2 infected hamsters revealed a primarily patchy and multifocal interstitial pneumonia (Fig. 3C) with lesion severity most prominent between 4–7 dpi. The histopathologic characterization of lesion spectra revealed (i) type II pneumocyte hyperplasia (Fig. 3D), (ii) interstitial and alveolar mixed mononuclear cell infiltrate (Figs. 3D-E), and (iii) perivascular edema and hemorrhage (Fig. 3E). Using scoring criteria developed by Gruber *et al*.^54^, the quantitation of the lung histopathology illustrated the progression of lung lesions in SARS-CoV-2-infected hamsters (Fig. 3F; 1±0, 4±2, 15±3 and 18±5 for mock, 2, 4 and 7 dpi, respectively), consistent with changes in L/BW ratio, gross pathology, and a previous report^53^. Importantly, these lesions have also been observed in COVID-19 patients^38,55,56^.

**Figure 3.**
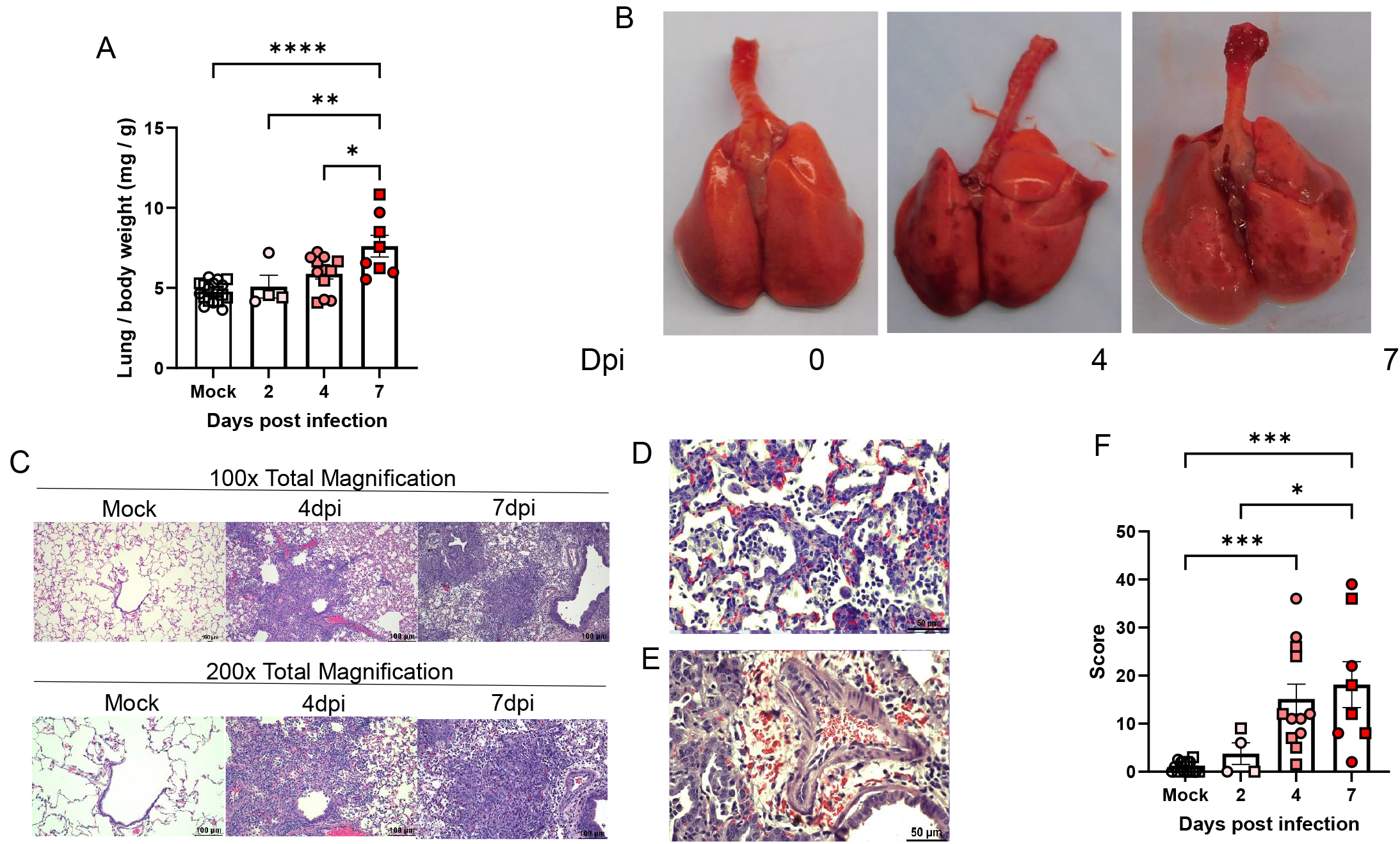
Lung injury after SARS-CoV-2 infection in hamsters. Golden Syrian hamsters were inoculated with SARS-CoV-2 as in Fig.1, then lung damage assessed by lung to body weight ratio (A), gross pathology (B), and histopathological analysis of the left lung, including representative hematoxylin and eosin (H&E) images (C-E) and quantitation by a blinded pathologist (F). (C) 100x (top) and 200x (bottom) total magnification of lungs from mock, 4dpi, and 7dpi displaying progression to interstitial pneumonia; 400x total magnification of (D) alveolus and alveolar interstitium displaying type II pneumocyte hyperplasia and mononuclear infiltrate and (E) a small-caliber artery displaying perivascular edema, hemorrhage, and intimal arteritis. Square is for males and circle for females. P<0.0001 (A and F) by one-way ANOVA, with *P<0.05, **P<0.01, ***P<0.001 and ****P<0.0001 by Tukey’s post-hoc test.

### SARS-CoV-2 causes defective mucociliary transport in hamsters

Having established that moderately severe SARS-CoV-2 respiratory infection in hamsters exhibit prominent airway involvement as with human COVID-19, we examined the effects of SARS-CoV-2 infection on the functional microanatomy of tracheas. Micro-optical coherence tomography (μOCT) allows the visualization and simultaneous quantitation of five microanatomic parameters that characterize the mucociliary clearance apparatus: airway surface liquid (ASL) depth and periciliary liquid (PCL) depth, tightly associated with ciliary height (CH), cilia beat frequency (CBF), the area of active ciliary beating, termed motile ciliation coverage (CC), and ultimately mucociliary transport (MCT) rate. The representative images of trachea for mock at 4 dpi, with corresponding m-mode resliced images to measure MCT rate of native particles within mucus, are shown in Figs. 4A and 4B (see Supplemental Videos 1 and 2). Multiple functional deficits were readily apparent, including diminished ciliary beating and reduced MCT rate upon SARS-CoV-2 infection. Particle transport quantification demonstrated the diminished MCT rate from 0.95±0.14 mm/min in mock to 0.31±0.17 mm/min (P=0.0479) and 0.36±0.10 mm/min (P=0.0076) at 2 and 4 dpi, respectively, whereas MCT had partially recovered by 7 dpi (0.72±0.16 mm/min; Fig 4C). This 67% and 62% decrease in MCT at peak infection (2 and 4 dpi, respectively) was closely associated with diminished motile CC, which was reduced from 23.9±2.8 in mock to 5.0±0.7 (P=0.0010), 6.6±1.3 (P<0.0001), and 14.1±2.7% (P=0.0324) at 2, 4 and 7 dpi, respectively, representing 21-28% of the normal level at peak infection that only partially recovered to 59% of mock by 7 dpi (Fig. 4D). Other notable features included an increase in ASL depth, likely reflecting the accumulation of mucus due to delayed transport rate at 4 dpi (from 9.0±1.4 to 15.8±2.0 μm, P=0.0100) that resolved as cilia began to recover at 7 dpi (9.4±0.5 μm *vs*. 4dpi, P=0.0420, Fig. 4E). PCL depth (Fig. 4F) exhibited no statistically significant changes, although a trend was evident, and measurements may have been limited by resolution under BSL3 containment. Residual cilia with detectable motion had no meaningful changes in mean CBF (Fig. 4G).

**Figure 4.**
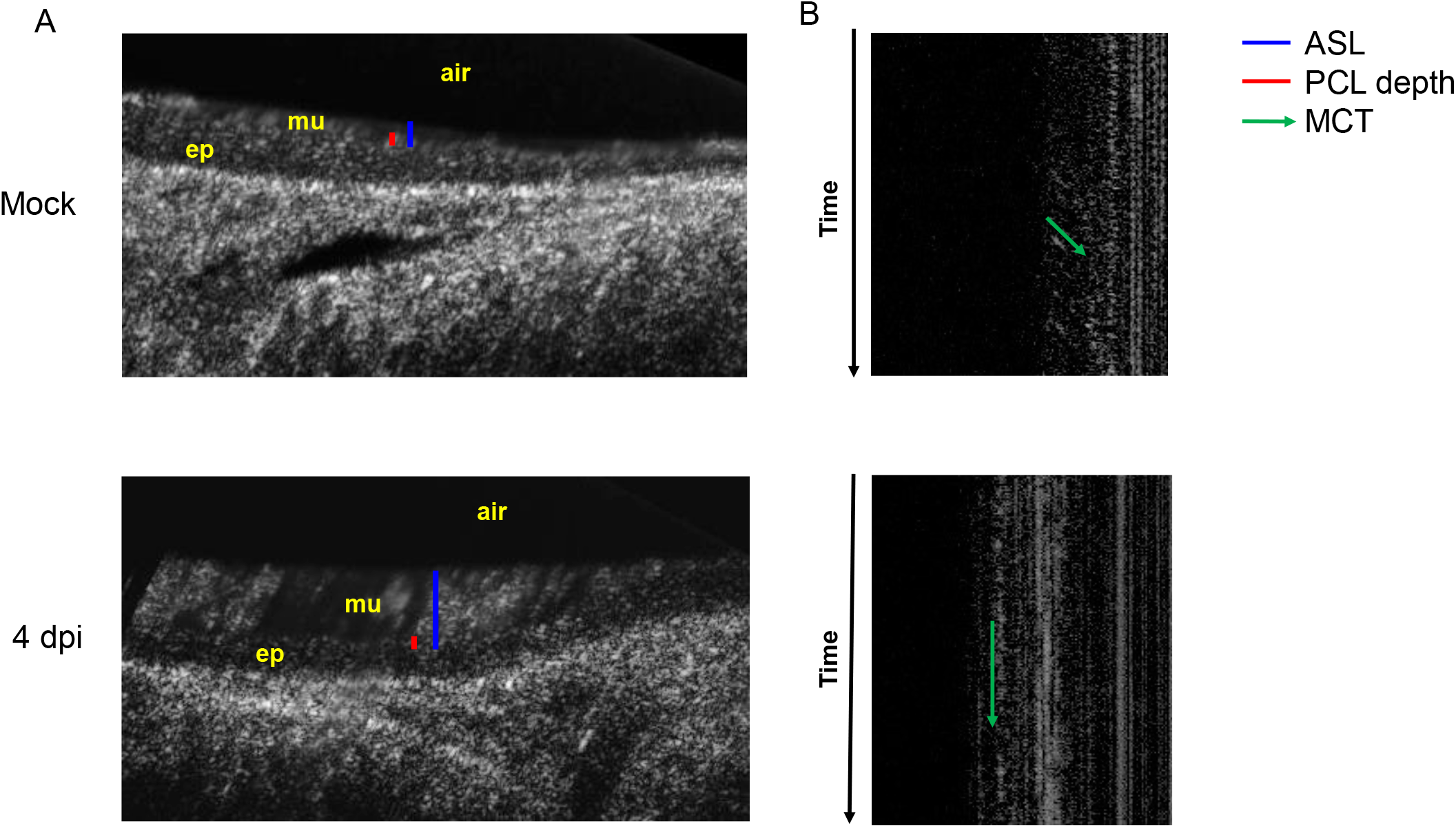
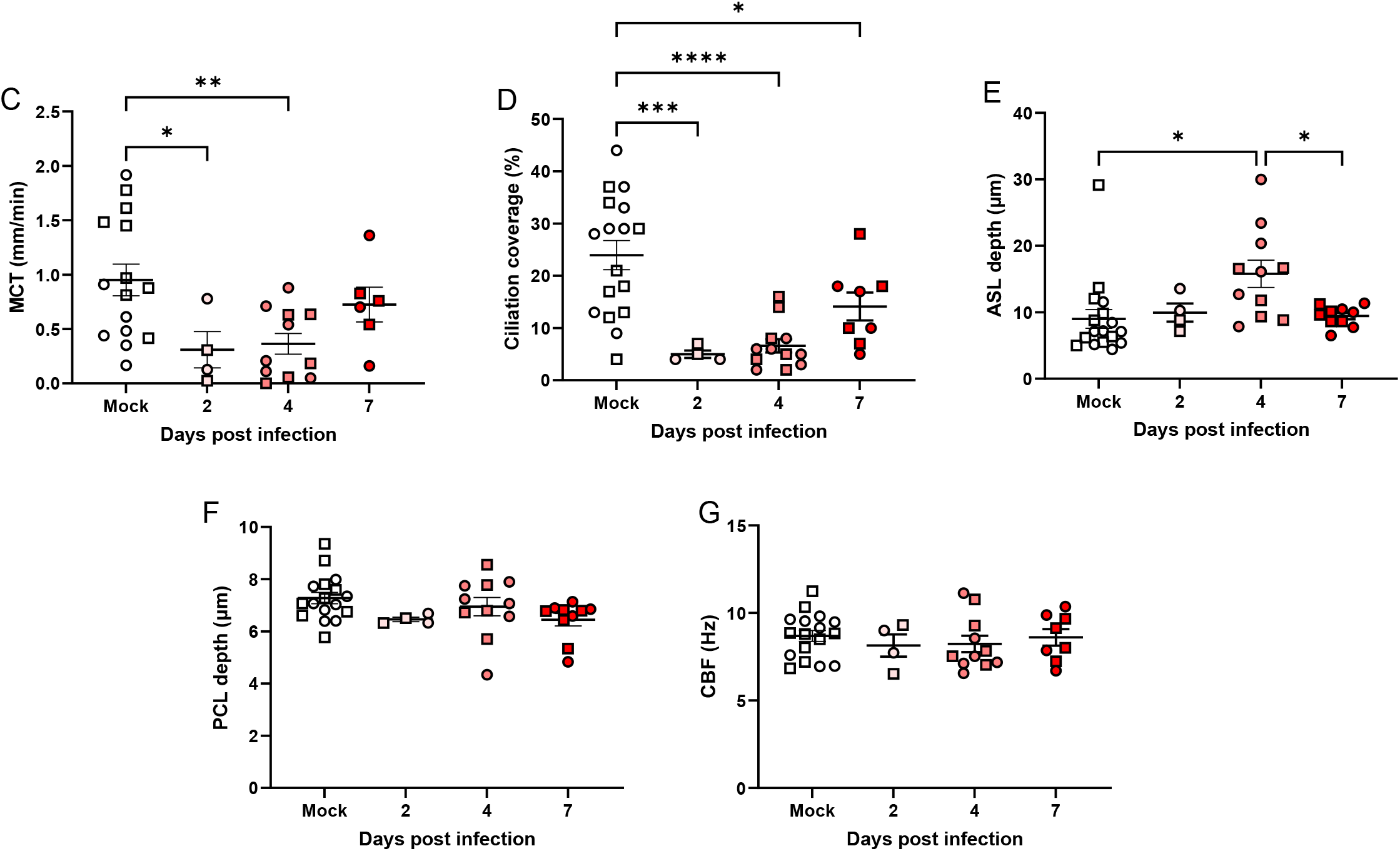
Mucociliary dysfunction in hamster airways after SARS-CoV-2 infection. Golden Syrian hamsters were inoculated in Fig.1 and following excision, tracheas were imaged by micro-optical coherence tomography (μOCT, A-G). Representative μOCT images from mock and 4 dpi (A) and M-mode projections of μOCT videos (B), ep: epithelium; mu: mucus. Mucociliary transport (MCT) rate (C), degree of active ciliation coverage (D), depths of airway surface liquid (ASL, E) and periciliary layer (PCL, F), and ciliary beat frequency (CBF, G) are shown. Square is for males and circle for females. P=0.0099, P<0.0001, P=0.0117, P=0.1106, P=0.7808 by one-way ANOVA for C-H, respectively; *P<0.05, **P<0.01, ***P<0.001 and ****P<0.0001 by Tukey’s post-hoc test.

To quantitatively ascertain contributors to the mucociliary transport defect due to SARS-CoV-2 infection, we performed regression analysis. The regression analysis of MCC parameters by animals demonstrated SARS-CoV-2 infection (ß=-0.495, P=0.003) and CC (ß=0.18, P=0.019) were each associated with delayed MCT (Table 1). To further explore the relationship between reduced MCT rate and SARS-CoV-2 infection, we conducted a linear mixed model analysis. Our initial model contained COVID status, dpi, and region of interest (ROI, 500mm specific field of view) as fixed effects, as well as a random animal-specific intercept. In this model, dpi and ROI was not statistically significant, but COVID status was statistically significant (P=0.0287). Consequently, ROI was removed as a fixed effect and replaced with sex and cohort as fixed effects with the random animal-specific intercept. This model, including experimental cohorts, dpi, sex, and infection status as fixed effects, confirmed that the presence of SARS-CoV-2 infection was the only variable that had significant effects on MCT rate (P=0.0389).

**Table 1:** Predictors of mucociliary transport burden by univariable regression. CH= ciliary height, ASL= airway surface liquid layer depth, CBF= ciliary beat frequency, CC= percentage of ciliary coverage

| Variable | $\beta$ | Standard Error | P-value | R<sup>2</sup> |
| --- | --- | --- | --- | --- |
| **COVID** | **-0.495** | **0.156** | **0.003** | **0.229** |
| CH | 0.119 | 0.087 | 0.181 | 0.052 |
| ASL | -0.014 | 0.014 | 0.347 | 0.026 |
| CBF | -0.052 | 0.066 | 0.436 | 0.019 |
| **CC** | **0.018** | **0.007** | **0.019** | **0.159** |

### SARS-CoV-2 causes tracheal and ciliary injury consistent with defective mucociliary transport in hamsters

Given prominent viral replication in the airway epithelia and defective MCT in hamster tracheas, we focused on the characterization of airway injury, an area recognized as increasingly important in the consequences of COVID-19 pathogenesis^19–21,28,29^. Representative images of H&E-stained tracheas (Fig. 5A) depicted normal pseudostratified and ciliated (apical surface) epithelium in mock hamster tracheas, while inflamed and barely ciliated epithelium in those at 4 dpi. Mononuclear inflammatory cells expanded the submucosa and infiltrated the epithelial mucosal layer (Fig. 5A). Cilia were largely absent, whereas apoptotic, desquamated epithelial cells that lost the attachment to adjacent epithelia were present (Fig. 5A). The injury was resolved at 7 dpi with recovered ciliated epithelia, and only occasional submucosal and intra-epithelial mononuclear cell infiltrate (Fig. 5A). Lesions in hamster tracheas were scored according to Gruber *et al*.^54^, and were summarized in Fig. 5B. The injury in the upper airways preceded that in the lung peaked at 2 dpi (5±0, P=0.0038 *vs*. mock) and 4 dpi (4±1, P<0.0001 *vs*. mock) and recovered by 7 dpi (2±0, P=0.0184 and 0.0007 *vs*. 2 and 4 dpi, respectively), consistent with the previous report^53^. This trend is correlated with the peak at 2 dpi and the subsequent decrease of viral titers in the hamster respiratory system.

**Figure 5.**
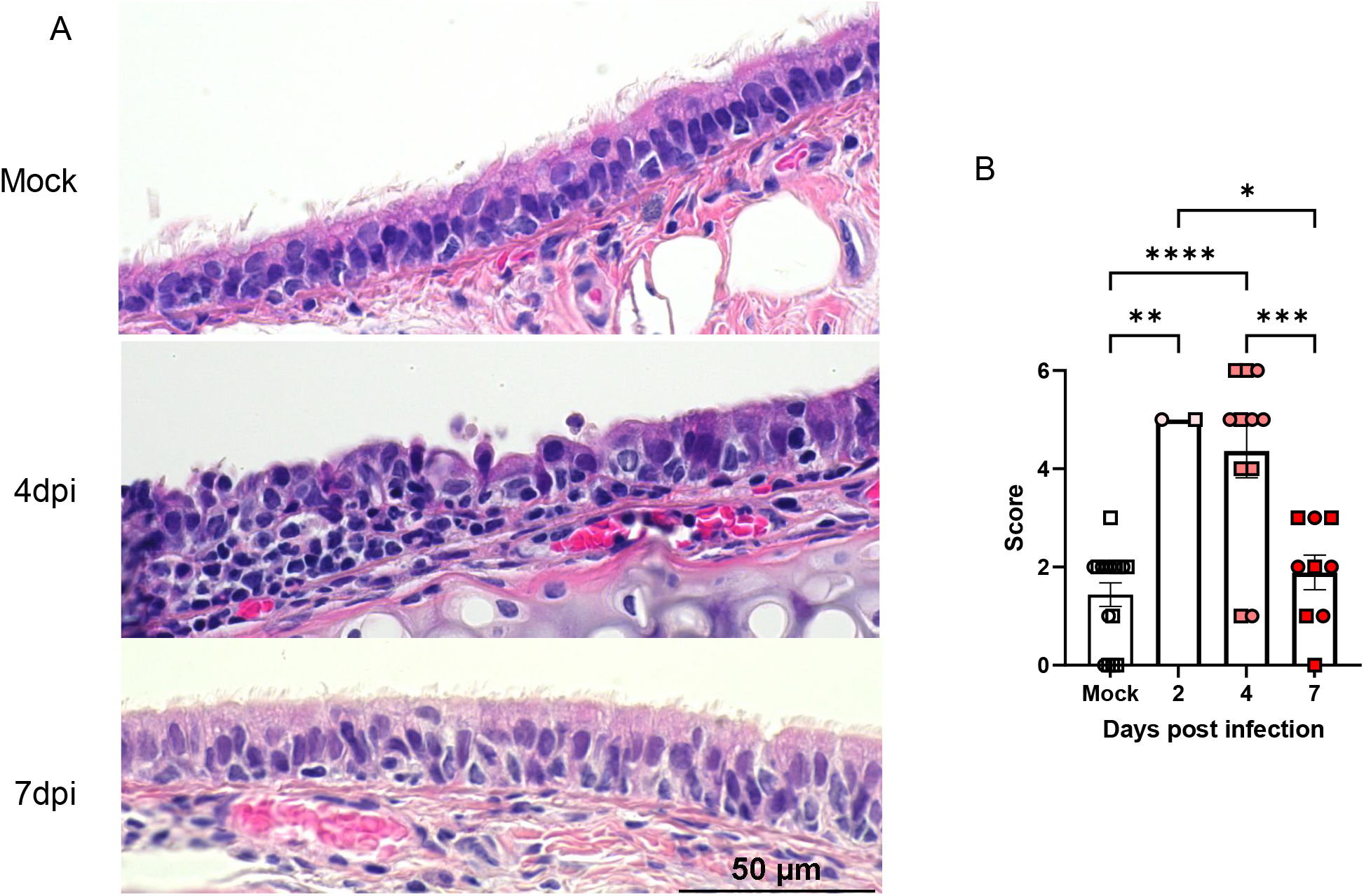
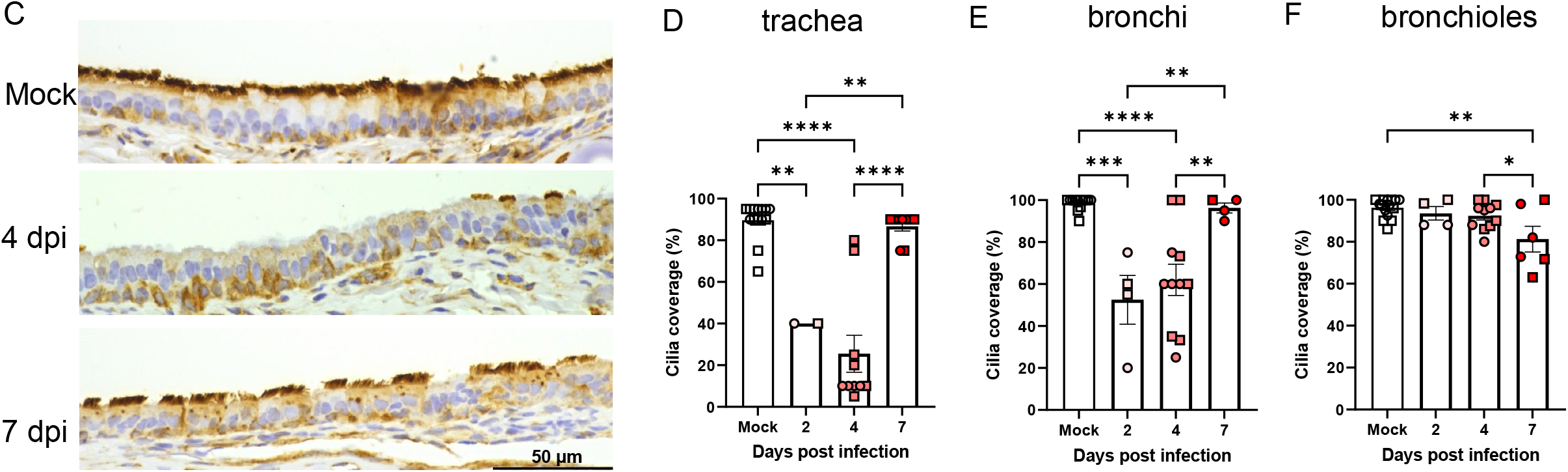
Tracheal injury after SARS-CoV-2 infection in hamsters. Golden Syrian hamsters were inoculated with SARS-CoV-2, then tracheas and lungs were processed for histopathological analysis. Representative H&E images (A) and quantitation by a blinded pathologist (B) are shown. Unstained slides were labeled with α tubulin by immunohistochemistry to specifically focus on ciliary injury. Representative images are shown of trachea (C), and quantitation of cilia coverage along the apical surface of the tracheal (D), bronchi (E), and bronchiolar (F) epithelia by a blinded investigator. Square is for males and circle for females. P<0.0001 (B and D) by one-way ANOVA, with *P<0.05, **P<0.01, ***P<0.001 and ****P<0.0001 by Tukey’s post-hoc test.

To examine the impact on cilia, we labeled unstained tracheal slides with α tubulin by immunohistochemistry (IHC). Tubulin positive layer along the apical surface of the tracheal epithelia dramatically decreased at 4 dpi compared to mock and then recovered at 7 dpi (Fig. 5C). Cilia coverage, the percentage of tubulin positive area along the ciliated airway epithelium, was quantified by a blinded pathologist (Fig. 5D). Results demonstrated cilia loss from a baseline of 90±2% in mock to 40±0% (P=0.0021) and 25±9% (P<0.0001) at 2 and 4 dpi (P=0.0055 and P<0.0001 *vs*. 2 and 4 dpi, respectively), whereas this was restored by 7 dpi, a finding in agreement with tracheal histopathological progression (Figs. 5A and B). This was also consistent with 95% loss of ciliated area found in hamster tracheas at 4 dpi with a different strain of SARS-CoV-2 (BetaCoV/France/IDF00372/2020)^29^. Examining both bronchi (Fig. 5E) and bronchiolar (Fig. 5F) compartments, the cilia expression of distal airways was also affected in a progressive fashion, although less severely. Notably, cilia loss has also been observed in other histological studies of SARS-CoV-2 infected hamsters^34,53^. More importantly, this time course of ciliary coverage changes is parallel with loss of motile CC (Fig. 4D) and MCT (Fig. 4C) measured by μOCT, indicating that the absence of cilia is the main contributor to the MCT deficit in SARS-CoV-2 infection.

### Aberrant ciliary motion and shortened residual cilia in COVID-19

Previously published data have established a direct correlation between CBF and MCT in the excised human^57^, porcine^58^, and ferret^59^ airways in the setting of diseases of mucus stasis such as cystic fibrosis and COPD. We assessed whether the relationship between MCT and CBF varied substantially by SARS-CoV-2 infection status when analyzed by each individual ROI. In mock controls, CBF positively correlated with MCT, whereas CBF negatively correlated with MCT in the presence of SARS-CoV-2 infection (r= −0.28, P=0.0390, Fig 6A). A sensitivity analysis examining the correlation of CBF with MCT limited to 4 dpi, when the severity of SARS-CoV-2 infection was at its peak, also revealed a negative correlation (r= −0.41, P=0.0064).

**Figure 6.**
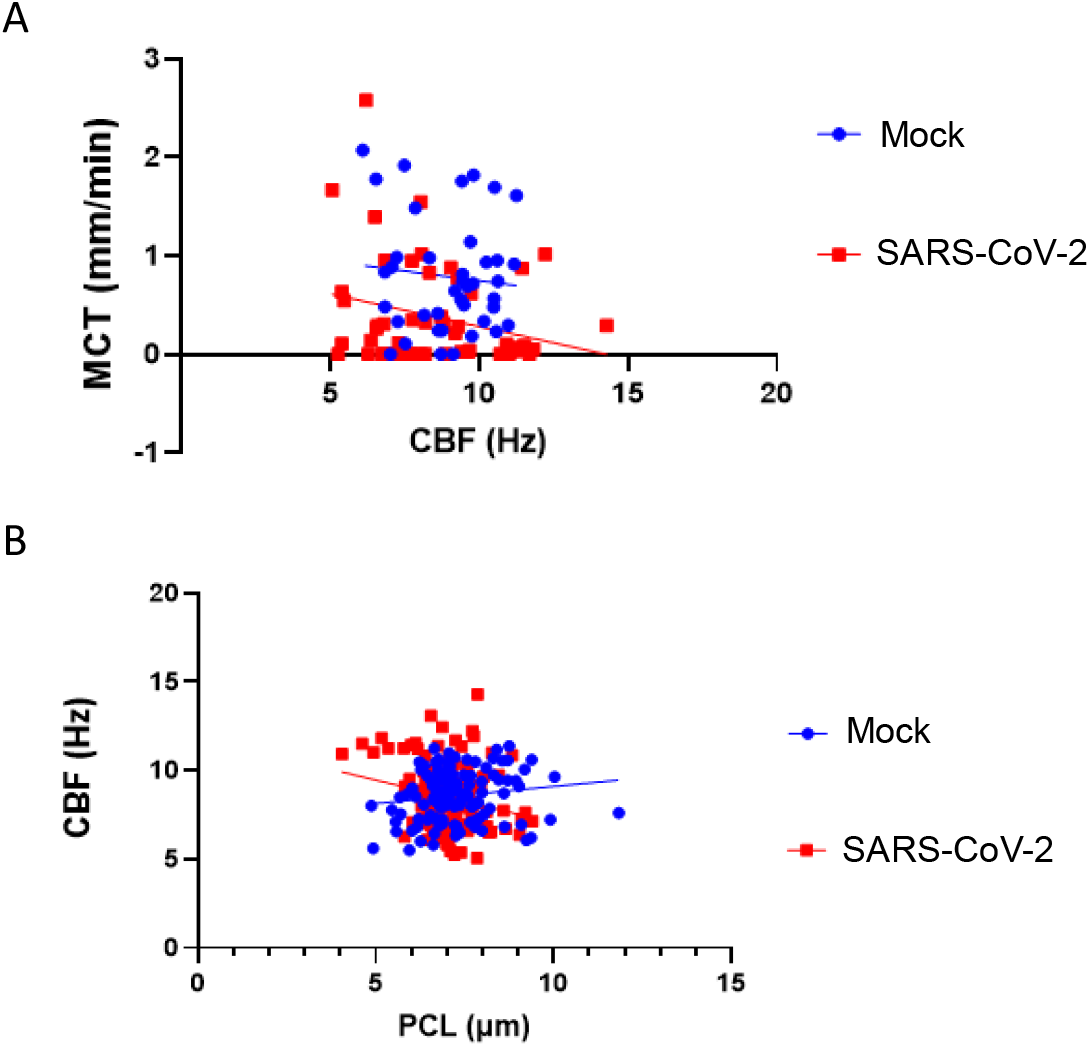
Association of MCC functional parameters with delayed MCT in SARS-CoV-2 infection. Linear correlation analysis of MCT with CBF (A); and PCL depth (B) in the presence and absence of SARS-CoV-2 infection. PCL: Periciliary liquid layer depth; MCT: Mucociliary transport rate; CBF: Ciliary beat frequency

Ciliary height has been reported to be reduced in SARS-CoV-2 infection in primary HBECs as part of the disordered repair associated with ciliated cell infection^29^. CBF is affected by the external environment in which they beat; shortened cilia would be expected to encounter less resistance from the overlying mucus gel, and thus potentially greater CBF for the same energy expenditure. This could explain the inverse relationship between CBF and MCT in SARS-CoV-2 infection. Indeed, we observed that PCL depth, a proxy for CH, was inversely associated with CBF, but only when SARS-CoV-2 infection was present (*r*= −0.12, P=0.0122; Fig. 6B). Thus, we suspect ciliary shortening detected at a regional level altered local ciliary beating, partially preserving CBF in these locations.

To examine ciliary function further, we examined the waveforms of individual cilia at 4 dpi as compared to mock controls (Fig. 7). The representative images of cilia maps showed fewer cilia with intact ciliary motion in SARS-CoV-2 infected hamsters (Fig. 7A) as compared to mock controls (Fig. 7A). The waveform analysis revealed preserved amplitude and consistent frequency in mock controls (Fig. 7B), whereas ciliary waveforms of SARS-CoV-2 infected hamsters exhibited variable amplitude and irregular beat patterns, even when mean CBF was maintained (Fig. 7B). Therefore, we conclude that fewer motile cilia combined with altered ciliary height and abnormal ciliary motion of residual cilia contribute to delayed MCT in SARS-CoV-2 infection.

**Figure 7.**
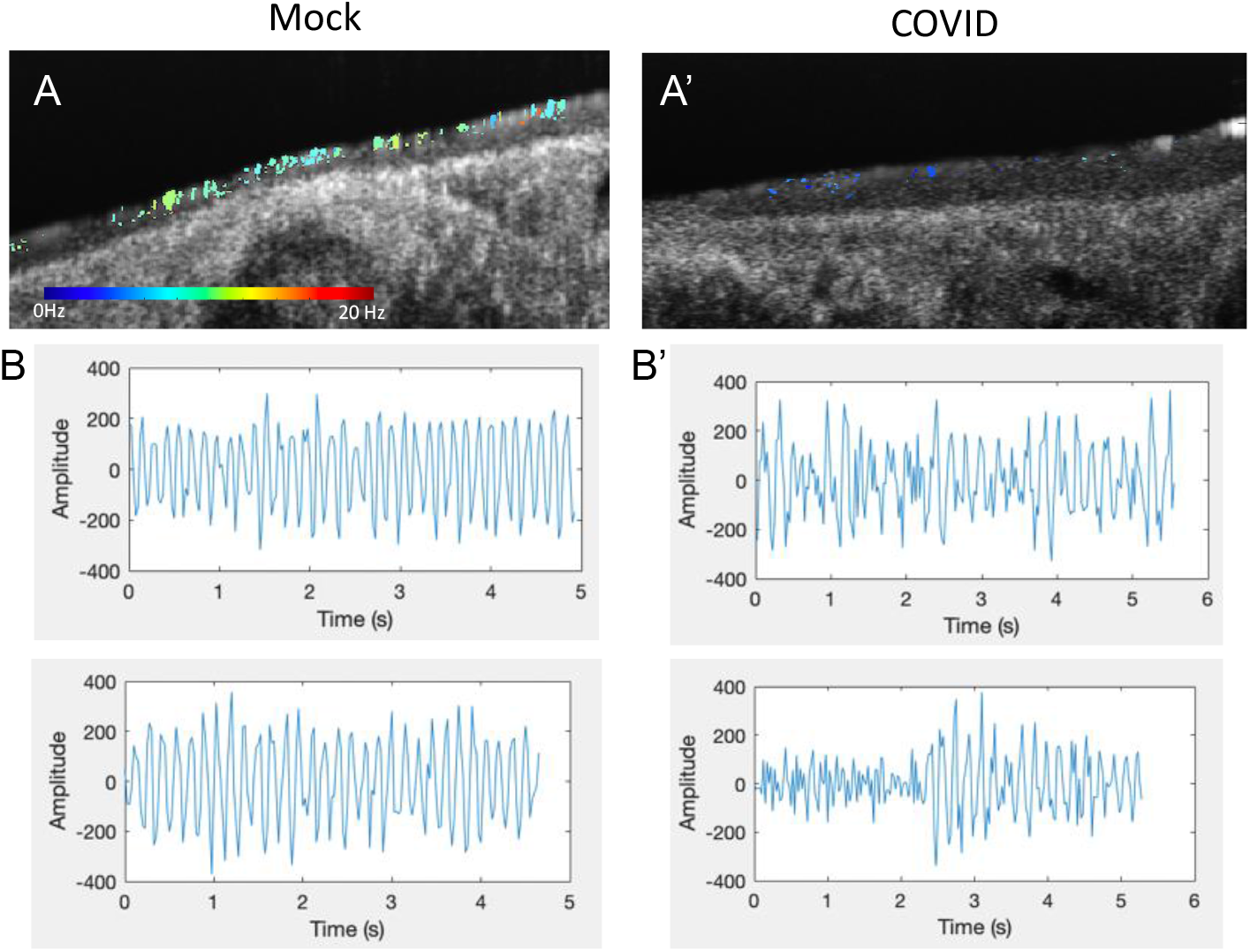
Abnormal ciliary beat frequency map and ciliary motion in SARS-CoV-2 infected hamsters. Compared to mock controls (A), fewer cilia with intact and maintained ciliary beat frequency was evident in SARS-CoV-2 infected hamsters (A’). Two representative waveform analysis of detected cilia exhibited consistent amplitude and frequency in trachea from mock controls (B) compared to erratic amplitude and irregular beat patterns in residual motile cilia in trachea from SARS-CoV-2 infected hamsters (B’).

## Discussion

Substantial clinical evidence supports the notion that ciliary function plays an important role in COVID-19 pathogenesis. Here, we quantitatively measured and comprehensively analyzed functional parameters of the MCT apparatus in Syrian hamsters, an animal model of COVID-19 pathogenesis, in parallel with virological and pathological analyses of SARS-CoV-2 infection. We noted evidence of steadily descending infection from the upper airways to the large and then small airways, as noted by markers of viral replication and consequently tissue injury. SARS-CoV-2 infection caused a 67% decrease in MCT as early as 2 dpi that was sustained through 7 dpi. Results corresponded with virological and pathological evidence of descending infection, primarily involving ciliated epithelial cells of the conducting airways, noting recovery of weight loss lagged behind. Corresponding with the functional deficit, we observed the loss of motile cilia, captured by reduced functional ciliary coverage, in addition to abnormal ciliary beating in residual cilia that corresponded with reduced MCT. The pathological analysis confirmed the deficit in ciliated respiratory cells. Of importance, some functional deficits in the MCC apparatus persisted through 7 days, as indicated by reduced CC, suggesting a propensity for secondary infection even after viral replication has ceased.

As in prior studies, we used functional analyses of the MCT apparatus to dissect relative contributors to the function decrement. While the cross-sectional analyses showed CBF was largely stable, the correlation analyses of co-localized areas provided additional insight. Results indicated the reduced ciliary length corresponded to greater CBF, an unexpected finding only observed in the presence of SARS-CoV-2 infection. We suspect this occurred by reducing ciliary workload and reducing cilia penetration into the mucus layer. Alternatively, it is also possible CBF was actively responding to the need to accelerate MCT through a feedback loop involving epithelial Ca^2+^ signaling, a mechanism known to occur when cilia encounter high workloads associated with increased mucus viscosity^60,61^. Cilia waveform analysis further suggested deranged ciliary beating and loss of coordinated motion. In total, our findings demonstrate that loss of cilia combined with the functionally inadequate beating of shortened cilia causes severe abnormalities in MCT.

MCT deficiency is likely to have significant clinical consequences. MCT decrements occurred early in the injury process, suggesting it may contribute to the propensity for descending viral infection, whereas parenchymal injury followed in time. The damage of the conducting airways appeared to result from direct cytopathic effects, as evidenced by detection of SARS-CoV-2 by in situ hybridization, whereas that of the parenchymal lung is likely due to downstream mediators, such as indirect injury and cytokine storm. The delayed MCT likely has other manifestations critical to COVID-19 pathogenesis. Mucus accumulation and stasis have been noted in the airways in autopsy studies of COVID-19^18–22^, and has contributed to complications in ICU studies^24–27^. In addition, mucus plugs reduce ventilation and alter gas exchange, which is likely to contribute to hypoxia in COVID-19 patients^20^. While our functional evaluation was limited to the trachea, we believe they are pertinent to the medium and small conducting airways. The altered function of the MCC apparatus would be expected to encourage secondary bacterial and fungal infections, as commonly associated with other diseases of mucus clearance, including post-acute influenza^62^, and increasingly reported as a late manifestation of COVID-19^24–27^. The functional deficit in functional CC observed in our study, despite resolution of ciliated cell loss by pathological analysis, suggests the potential to contribute to persistent respiratory manifestations. Aspergillus infection is highly associated with abnormal mucus clearance in other diseases^63^ and thus may explain its increasingly recognized role as a COVID-19 complication, along with changes in immunity as a response to viral infection^64–67^.

The degree of body weight loss post-infection is comparable to previous hamster models regardless of the differences in the viral strain and inoculation titers^34,37–39,41,42,44,46^. The preferential viral infection in the respiratory tract^9^, the propensity for descending infection, and the specific lesions in the lung have been observed in human autopsy^38,55,56^. A 95% loss of ciliated area has been found in hamster tracheas at 4 dpi with a different strain of SARS-CoV-2^29^, consistent with 72% loss of cilia coverage by histopathology at 4 dpi observed here. The cilia damage post-infection has been demonstrated in HTBE cells^28,29^ and hamster tracheas^34,53^. The consequences have been detected by decreased speed and non-directional movement visualized in HTBE cells by polystyrene beads^29^. While we did not investigate cellular mechanisms of ciliated cell loss, downregulation of Foxj1, the regulator for ciliogenesis, has been suggested as the upstream mechanism of cilia loss in SARS-CoV-2 infected HTBE cells immediately post-infection^29^. The loss of ciliated cells is likely to explain scRNA-seq analysis of bronchial alveolar lavage fluid (BALF) from severe COVID-19 patients showing down-regulation of gene transcripts for ciliary structure and function^19^. Further, MCT deficiency could contribute to elevated protein levels of mucin in the airways, the detection of mucin mRNA in blood, and abnormal mucus viscosity detected in severe COVID-19 patients^19,21,68^.

In summary, we detected and quantitated MCT deficiency in an animal model of COVID-19. By comparing MCT trajectory and time courses of physiological, virological, and pathological progression in hamsters infected with SARS-CoV-2, we suggest that functional deficits of the MCT apparatus predispose to COVID-19 pathogenesis by extending viral retention and is a risk factor for secondary infection. Monitoring abnormal ciliated cell function may indicate progression and represent a treatment opportunity for COVID-19. Furthermore, our results strongly suggest that additional therapies directed towards the MCT apparatus deserve further investigation as a treatment modality.

## Methods

### SARS-CoV-2

The SARS-CoV-2 isolate USA-WA1/2020 was obtained from BEI Resources (cat. # NR-52281, Manassas, VA) and further propagated in Vero E6 cells (cat. # C1008, ATCC) through two more passages to obtain a working stock of virus at sufficient titer. Viral titers were determined by standard plaque assay^69^. Briefly, virus was serially diluted and added to confluent Vero E6 cells grown on 6-well plates. After one-hour incubation, 1X Minimal Essential Medium (Gibco^™^ # 11430-030 from Fisher) with 2% fetal bovine serum (cat. # C788U20 from Denville Scientific, Inc.), 1X antibiotic-antimycotics (Gibco^™^ # 15240-062 from Fisher), and 0.6 % Avicel CL-611 (from FMC BioPolymer, Philadelphia, PA) was added, and the infection was allowed to proceed for 72 hr. Plates were fixed by submerging in 10% phosphate-buffered formalin (cat. # SF100-20 from Fisher) for 24 h, stained with 0.05% neutral red (cat. # N2889-100ML from Sigma) for about 1 hour, and rinsed in water. Plaques were counted manually from the scanned image of each well. RNA Sequencing by UAB Heflin Genomics Core Laboratory indicated less than 10 mutations from the parent strain that is no more than 14% of the sequences annotated. The aliquots of viruses were stored at −80°C for up to 3 months prior to usage. The viral transport medium (VTM) containing Hanks Balanced Salt Solution [Thermo Fisher Scientific (Fisher) cat. # 14-025-092], 2% heat-inactivated fetal bovine serum (Fisher cat. # 26-140-079), 100 μg/mL gentamicin (Sigma cat. # G1397) and 0.5 μg/mL amphotericin B (Sigma cat. # A2942) were prepared according to the CDC Standard Operating Procedure (SOP# DSR-052-05)^70^.

### Animal model and sample collection

All procedures involving animals and SARS-CoV-2 were approved by the Institutional Animal Care and the Use Committee (# 20232) and Institutional Biosafety Committee (# 10-262) at the University of Alabama at Birmingham (UAB). LVG Golden Syrian hamsters were purchased from Charles River (cat. # Crl: LVG(SYR)/049) and were inbred in the animal facility at UAB. Sex-matched adult hamsters were inoculated intranasally (50 μl per nare) with 3 × 10^5^ plaque-forming units of SARS-CoV-2 or vehicle (mock). Clinical signs including body weight, lethargy, fur, posture, sneezing, coughing, diarrhea, and nasal or ocular discharge were monitored; nasal brushes, oral and rectal swabs were collected; and hamsters were euthanized up to seven days post-infection. After euthanasia by intraperitoneal injection of 100 mg/kg pentobarbital, the ventral side of tracheas were excised and prepared for μOCT imaging followed by fixation in 10% neutral buffered formalin (NBF, Fisher cat. # 23-245684) for histological and immunohistochemistry (IHC) analysis; lungs were cleaned and weighed; blood, nasal washes and bronchial alveolar lavage fluid (BALF) from right lung lobes were collected for quantitation of both genomic and subgenomic viral RNA by qRT^2^-PCR; and left lungs were slowly inflated with 3 ml of NBF and then fixed in NBF for histological and RNAscope^®^ analysis. Tracheas and left lungs were fixed in NBF for 3-5 days to assure viral deactivation before being processed by the UAB Comparative Pathology Laboratory.

### Viral detection by quantitative real-time polymerase chain reaction (qRT^2^-PCR)

Using GUM - 877A Proxabrush from Amazon, nasal brushes were collected, and oral and rectal swabs were collected using Medical Packaging Swab-Pak^™^ Swabs from Fisher (cat. # 22-281661). Once collected, tips of brushes and swabs containing samples were placed in 0.5 ml VTM and mixed well. Blood was collected into BD Microtainer^™^ Capillary Blood Collector (Fisher cat. # 02-675-185) through the cardiac puncture. Serum was isolated within 10 min of collection to avoid hemolysis and through centrifugation at 1,000 g for 10 min. For nasal washes, the laryngeal or proximal trachea was clamped by hemostats, 0.5 ml VTM was injected into one nostril through a blunted needle, and the liquid eluting out of both nostrils were collected, the air was then injected to collect the residual liquid in nasal turbinate. BALF was collected from right lung lobes using 2ml of PBS (pH 7.4, Fisher cat. # 10-010-023) with 60-80% return.

Maxwell^®^ RSC Viral Total Nucleic Acid Purification Kit (cat. # ASB1330, Promega Corporation, Madison, WI) was used for RNA isolation. As instructed by the manufacturer, 300μl of sample was mixed by vortexing for 10 sec with 330 μl of a freshly made mixture of lysis buffer and proteinase K solution at a 10:1 ratio. Upon incubation at room temperature for 10min and at 56°C for additional10min, RNA was extracted utilizing the Maxwell^®^ RSC 48 Instrument (Promega Corporation), and qRT^2^-PCR was performed utilizing QuantStudio^™^ 5 Real-Time PCR instrument (Fisher). The genomic RNA PCR primers (forward 5’-GAC CCC AAA ATC AGC GAA AT-3’ and reverse 5’-TCT GGT TAC TAC TGC CAG TTG AAT CTG-3’) and probe (/SFAM/ACC CCG CAT TAC GTT TGG TGG ACC/BHQ_1) targeted the nucleocapsid (N) gene and were purchased from Integrated DNA Technologies, Inc. (IDT, Coralville, Iowa). AccuPlex^™^ SARS-CoV-2 Reference Material (cat. # 0505-0126, SeraCare Life Sciences, Inc., Milford, MA) was extracted and amplified in parallel to generate a standard curve enabling viral quantitation. The subgenomic (sg) PCR assay utilized a forward primer targeting the leader sequence (5’-CGA TCT CTT GTA GAT CTG TTC TC-3’, from IDT) and the same reverse primer targeting the N gene as for genomic RNA PCR. The PCR products were detected by *Power* SYBR^™^ Green RNA-to-C-^™^ *1-Step* Kit (Fisher cat. # 4389986). The sgRNA standard was constructed in-house by amplifying the N gene using primers sgRNA-UF (5’-CCA ACC AAC TTT CGA TCT CTT GTA-3’) and sgRNA-NR1 (5’-AAG GTC TTC CTT GCC ATG TTG-3’) from IDT. The amplified band was purified from agarose gel, and this stock standard was quantitated as 2×10^13^ copies/ml by NanoDrop^™^ One Microvolume UV-Vis Spectrophotometer (Fisher), which was serially diluted and utilized directly as the PCR template to generate standard curve for viral sgRNA quantitation.

### Viral detection by RNAscope^®^

The viral RNA in the left lung was detected using RNAscope^®^ 2.5 HD Reagent Kit - RED (cat. # 322350 from Advanced Cell Diagnostics, Inc., Newark, CA) and following manufacture instructions. Briefly, unstained lung slides were deparaffinized and pretreated with hydrogen peroxide, target retrieval reagent, and then Protease Plus. The viral RNA was hybridized to a probe specific to USA-WA1/2020 strain, V-SARS-CoV-2-N-O1 (cat. # 863831, Advanced Cell Diagnostics, Inc., Newark, CA). Following amplification, RED solution was added to visualize the signal, and slides were counterstained and then mounted. In parallel, a probe that hybridizes gene of a common protein peptidylprolyl isomerase B (PPIB) and is specific to a hamster, Mau-Ppib (cat. # 890851, Advanced Cell Diagnostics, Inc., Newark, CA), was applied to positive control slides, and a probe for a soil bacterial gene, DapB (cat. # 320871, Advanced Cell Diagnostics, Inc., Newark, CA), to negative control slides.

### Histological analysis

H&E stained trachea and lung slices were blinded and semiquantitatively scored by a board-certified veterinary anatomic pathologist following published criteria^54^. Briefly, tracheal, alveolar, and interstitial (AI), bronchiolar and peribronchiolar (BP), and vascular (V) lesions were scored based on subcategories such as necrosis, cell debris, and location-specific criteria with the presence of these lesions = 1 and their absence = 0. The subcategory scores were added up as scores for these sublocations. For the lung, the sum of sublocation scores (AI + BP + V) were multiplied by overall extent of affected area (0 – 4 with 0 = normal, 1 = 10 – 25%, 2 = 25 – 50%, 3 = 50 – 75%, and 4 > 75% of lung surface area displaying pathologic lesion spectra, respectively) of the whole lung slice to generate a final histopathological score of the left lung.

### Immunohistochemistry (IHC) staining and analysis

Unstained tracheal slides were labeled with α tubulin by UAB Pathology Core Research Laboratory. Standard IHC process was applied with antigen retrieval by 0.01 M sodium citrate buffer (pH 6), 1:30,000 dilution of anti-acetylated tubulin (cat. # T7451 from Sigma-Aldrich) and 1:500 dilution of goat anti-mouse IgG H&L secondary antibody conjugated with HRP (Abcam ab6789). The percentage of tubulin positive signal (cilia presence) along the apical surface of the tracheal epithelia (cilia coverage) were semi quantitated by a blinded pathologist.

### Micro-optical coherence tomography (μOCT)

Excised trachea was imaged from hamsters, as previously described for other species^57–60^. Once the larynx area was open, and the trachea was exposed under euthanasia, connective tissues around the trachea were removed, ventral side of tracheas was excised, immediately placed and maintained on a gauze saturated with room temperature Dulbecco’s Modified Eagle Medium (Fisher cat. # 11-965-092). The trachea was sealed using Parafilm (Fisher cat. # 50-136-7664) in a petri-dish with an assembled window made of a scratch- and UV-resistant acrylic sheet of 1/16” thickness (cat. # 1227T739 from McMaster-Carr, Princeton, NJ) to preserve laser transmission and thus imaging quality, by μOCT. The instrumentation, methodology, and analysis of μOCT were performed as previously established^52,57,58^ and applied to both animal and human tracheas^57–60^. Up to nine regions of interest per hamster trachea were imaged. Airway surface liquid (ASL) depth, periciliary layer (PCL) height, cilia beat frequency (CBF), mucociliary transport (MCT) rate, and ciliation coverage (CC) on epithelial surfaces of tracheas were analyzed using ImageJ (NIH, Bethesda, MD) and MATLAB software (MathWorks, Natick, MA)^57–60^. The peak frequency described ciliary beat frequency in the power spectrum. A Fourier power spectrum was computed for 3×3-pixel sub-regions in the μOCT videos along the motile cilia interface. Low-frequency vibrations were removed by applying a temporally high pass filter with a cut-off frequency of 3 Hz to negate the confounding signal. The resulting 20 sub-regions with the highest frequency peak sharpness, defined as the peak power density divided by the total power density, were then analyzed to generate mean CBF for each ROI.^52^ The CBF map was generated through a MATLAB script to visualize the location and frequency of active cilia across the epithelial cell surface and the proportion of pixels with active cilia quantified as previously described^71^. To characterize ciliary waveforms of detected cilia, the amplitude of the ciliary signal generated by ciliary motion was plotted for selected sub-regions of interest characteristic of mean values^71^.

### Statistical analysis

Data were analyzed using unpaired t-test, one-way or two-way analysis of variance (ANOVA) followed by recommended Tukey’s or Šídák’s multiple-comparison tests, descriptive statistics, and regression methods (both univariable regression and linear mixed models) in GraphPad Prism 9.2.0 (GraphPad Software, Inc., La Jolla, CA), SPSS Statistics (IBM, Armonk, NY), or SAS v9.4 (Cary, NC). Data are presented as mean ± standard error, with p-values < 0.05 considered statistically significant. Correlations between metrics were calculated using linear or semi-log regression methods. A univariable linear regression model was performed to assess for effects of ASL, CH, CC, and CBF on mean MCT. A linear mixed model for repeated measures analysis was used to model MCT as a repeated measure across ROI (technical and biological replicates), and fixed effect predictors included ROI effects, cohort effects, sex effects, infection effects, and the potential for interaction among them. Random effects included random intercept only. The final models omitted interaction terms between predictor variables and ROI or cohort, as these were not statistically significant and did not alter conclusions.

## Acknowledgements

We thank UAB Comparative Pathology Laboratory for tissue processing, slicing and H&E staining, and UAB Pathology Core Research Laboratory for α-tubulin IHC staining. This work was supported by: R35 HL135816-04S1 (Rowe), P30DK072482-12 (Rowe), 5F31HL146083-02 (Lever), 3T32GM008361-30S1 (Lever), 2T32HL105346-11A1 (Lever), and Cystic Fibrosis Foundation-PHILLI20G0 (Philips/Campos-Gómez).

